# Noisy metabolism can drive the evolution of microbial cross-feeding

**DOI:** 10.1101/2021.06.02.446805

**Authors:** Jaime G. Lopez, Ned S. Wingreen

## Abstract

Cross-feeding, the exchange of nutrients between organisms, is ubiquitous in microbial communities. Despite its importance in natural and engineered microbial systems, our understanding of how cross-feeding arises is incomplete, with existing theories limited to specific scenarios. Here, we introduce a novel theory for the evolution of cross-feeding, which we term noise-averaging cooperation (NAC). NAC is based on the idea that, due to their small size, bacteria are prone to noisy regulation of metabolism which limits their growth rate. To compensate, related bacteria can share metabolites with each other to “average out” noise and improve their collective growth. This metabolite sharing among kin then allows for the evolution of metabolic interdependencies via gene deletions (this can be viewed as a generalization of the Black Queen Hypothesis). We first characterize NAC in a simple model of cell metabolism, showing that metabolite leakage can in principle substantially increase growth rate in a community context. Next, we develop a generalized framework for estimating the potential benefits of NAC among real bacteria. Using single-cell protein abundance data, we predict that bacteria suffer from substantial noise-driven growth inefficiencies, and may therefore benefit from NAC. Finally, we review existing evidence for NAC and outline potential experimental approaches to detect NAC in microbial communities.

## Introduction

Microbial communities are found nearly everywhere in nature, inhabiting ecosystems ranging from hydrothermal vents [1] to mammalian guts [2]. One of the most striking properties of these communities is the ubiquity of cooperation: microbes frequently share resources. This exchange of resources is broadly referred to as ‘cross-feeding’. Cross-feeding is widespread across both natural and engineered systems, with notable examples occurring in the human gut and in wastewater treatment systems [3, 4]. These metabolic interactions link organisms across the entire tree of life, occurring both within and between kingdoms [5] and even between specialized microbes of the same species [6].

Cross-feeding plays a major role in the structure and function of microbial communities. In natural settings, cross-feeding is known to be a significant driver of microbial diversity, allowing many species to coexist on a small number of primary resources [7, 8]. This microbial diversity has been linked to a wide variety of community properties [9, 10], including influence on host fitness [11]. Cross-feeding can even play a role in public health: it has been shown that metabolite exchange can allow pathogens to compensate for fitness losses associated with antibiotic resistance [12]. In engineered systems, cross-feeding can be necessary for efficient operation. For example, wastewater treatment reactors rely on cross-feeding to prevent the buildup of inhibitory waste products [13]. Thus, a thorough accounting of the factors promoting cross-feeding is an important part of understanding both natural and engineered microbial communities.

As a result of cross-feeding’s key role in microbial communities, much work has been dedicated to unraveling its evolutionary origin. There are broadly two different types of cross-feeding, each speculated to have its own mechanisms of evolution. ‘Waste-product cross-feeding’, in which an organism feeds on the waste products of another organism, is theorized to evolve as a result of growth trade-offs that make it optimal for organisms to only partially metabolize substrates. This partial metabolism leads to the secretion of compounds that can be further metabolized by downstream organisms [14–16]. (2) In contrast, ‘metabolite cross-feeding’, in which organisms share metabolites that they themselves require, is often explained by invoking the Black Queen Hypothesis (BQH) [17–20]. This hypothesis posits that that if a function is ‘leaky’ (i.e. can benefit organisms not performing the function), there will be a selective advantage for some organisms to lose the function and rely on leakage from others. This type of gene loss can ultimately lead to a web of interdependencies.

While the BQH provides a plausible mechanism for the evolution of some cross-feeding relationships, its underlying assumption that many metabolites are naturally leaky is not well-supported. For some functions, leakage is clearly unavoidable because key processes take place outside the cell, such as iron uptake via siderophores or hydrolysis of large polymers by extracellular enzymes [21, 22]. However, many cross-feeding relationships involve metabolites that are produced intracellularly, and it is generally not known how these metabolites exit the cell, much less that this leakage is inevitable. Polar or charged metabolites are known to have low membrane permeability, limiting the possibility of natural leakage through the cell membrane [23]. Indeed, even if a metabolite has a high membrane permeability, this does not necessarily indicate high absolute leakage rates. Cells could minimize leakage by maintaining only a small metabolite pool with rapid turn-over or by storing the metabolite in an altered, less leakage-prone form. Even if substantial quantities of a metabolite are observed to escape via a leaky membrane, it cannot be determined without further study whether this leakage is truly inevitable or, rather, advantageous in some way. Thus, there is motivation for a more general theory of cross-feeding that can better explain the origins of leakiness. Can cross-feeding evolve for metabolites that are not naturally leaky?

Here, we explore a novel mechanism for the evolution of cross-feeding in microbial communities. The basis of this mechanism is that metabolic enzyme regulation is inevitably imperfect, and particularly so in bacteria due to their small size. The resulting imbalances in enzyme levels can lead to growth inefficiencies due to the under or overproduction of necessary metabolites. Rather than attempting to downregulate the activity of the excess enzymes, cells can in principle improve their communal growth rate by exchanging metabolites among their kin and effectively “averaging” out intracellular noise. We term this mechanism noise-averaging cooperation (NAC). We first characterize NAC in a model of a small population of cells, demonstrating that metabolite leakage can increase collective fitness. We find that in extreme cases, NAC can even prevent the death of cells whose poor enzyme regulation would otherwise lead to irreversible growth arrest. We then develop a generalized, experimentally accessible framework for estimating how NAC is influenced by community size and the complexity of metabolic pathways. Using this framework and single-molecule data on *Escherichia coli* enzyme levels, we predict that typical bacteria suffer from significant growth inefficiencies due to imperfect regulation, and thus could benefit from metabolite exchange. In turn, the evolution of beneficial metabolite exchange among kin creates the conditions for interdependencies to evolve via the BQH and gene deletions. Thus, the proposed mechanism provides a more general theory of cross-feeding evolution in which metabolic leakiness is not assumed *a priori*, but rather arises from evolutionary pressures.

## Results

### A. Isolated cells

We begin by exploring the impact of enzyme noise on the growth of an isolated model cell. To focus on the role of enzyme level fluctuations, we consider a cell of fixed volume and track the numbers of internal enzyme and metabolite molecules. Cell growth rate is recorded, but does not explicitly lead to an increase in cell volume. Instead, to capture the effects of cell growth and the associated volume increase, the rates of enzyme production and enzyme loss by dilution are both taken to be proportional to the growth rate. Since metabolite production and consumption fluxes are generally large compared to the dilution of metabolite levels by growth, we neglect the small effect of dilution on the metabolite levels. In this simple model, we assume that cell growth requires two essential metabolites that have intracellular counts of 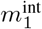 and 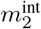. These metabolites are produced intracellularly by specialized enzymes with intracellular counts of *E*_1_ and *E*_2_. Both metabolites are produced from the same precursor, which is imported such that a constant number of precursor molecules is maintained within the cell. Each enzyme produces its metabolite at rate *κE_i_*, with *κ* encompassing both the precursor concentration and the enzyme rate constant. In accordance with Liebig’s law of the minimum, growth is proportional to the level of the least abundant of the two metabolites such that the growth rate is 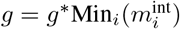, where *g*^∗^ is a constant relating the metabolite levels to the cell growth rate. Thus, the cell growth rate is maximized when the metabolites are produced and present in equal amounts. Metabolites within the cell are assumed to be consumed at a rate proportional to the growth rate. Metabolites are also exchanged with an extracellular space whose volume is *r*_V_ times greater than a cell volume. This exchange occurs via membrane diffusion with permeability *P* (we show in Appendix A that this form of exchange is mathematically equivalent to active transport in the linear regime). Metabolites both inside and outside the cell are passively degraded at a rate *δ*. A schematic of this model is shown in Figure 1A. Formally, the intra- and extracellular counts of the metabolites evolve according to the following nondimensionalized equations (see Appendix A for details):

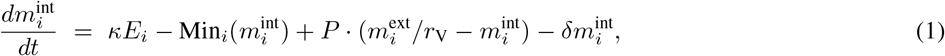

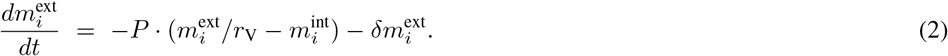

We model these metabolite dynamics as deterministic, as there is generally a large number of each essential metabolite within cells [24].

**FIG. 1:**
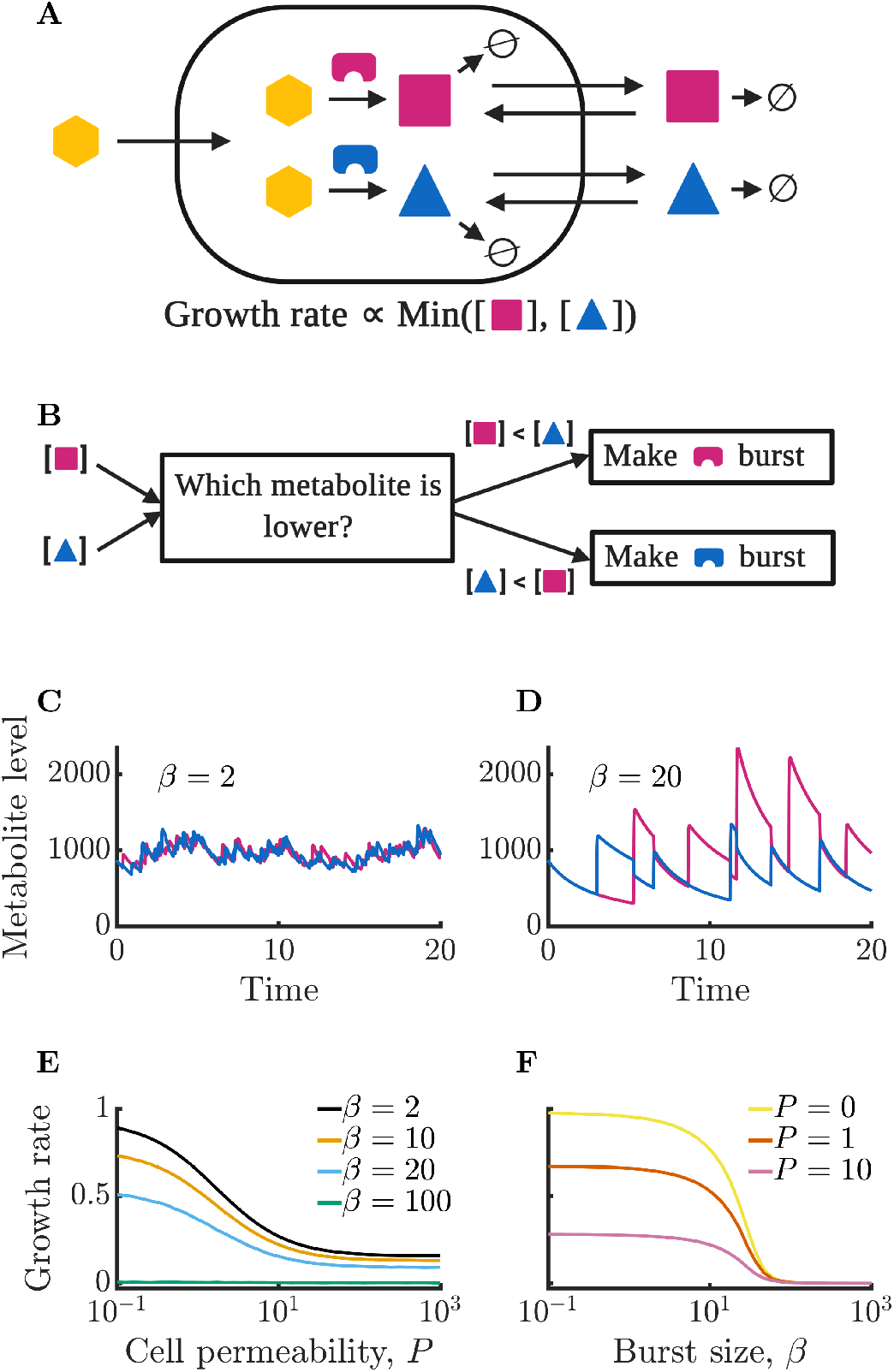
Isolated bacterial cells suffer negative growth effects from noisy enzyme regulation and metabolite leakage. (*A*) Schematic of modeled intracellular dynamics. Cells import an external nutrient (yellow hexagon) that can be converted by enzymes (magenta and blue) into two essential metabolites. Metabolites passively exchange with the extracellular medium (“leakage”), and degrade at a fixed r ate. The two metabolites are used for growth in accord with Liebig’s law of the minimum. (*B*) Schematic of dynamic enzyme regulation scheme: the type of enzyme produced is always the one associated with the lower metabolite pool. (*C*) Metabolite timecourse of a cell that produces enzymes in small bursts. See Eqs. 1–2 for details; parameters *β* = 2, *γ* = 50, *κ* = 100, *r*_V_ = 10, *P* = 0.3, *δ* = 1, *g*^∗^ = 1 10^−5^. (*D*) Metabolite timecourse of a cell that produces enzymes in large bursts. Same parameters as in *C* but with *β* = 20. (*E*) Average growth rate of an isolated cell for differing values of permeability; parameters as in *C* and *D* except as specified. Growth rate is normalized to the maximum possible growth rate,i.e. with perfect regulation and zero permeability. (*F*) Average growth rate of an isolated cell for differing values of burst size. Parameters as in *C* and *D* except as specified.

**FIG. 2:**
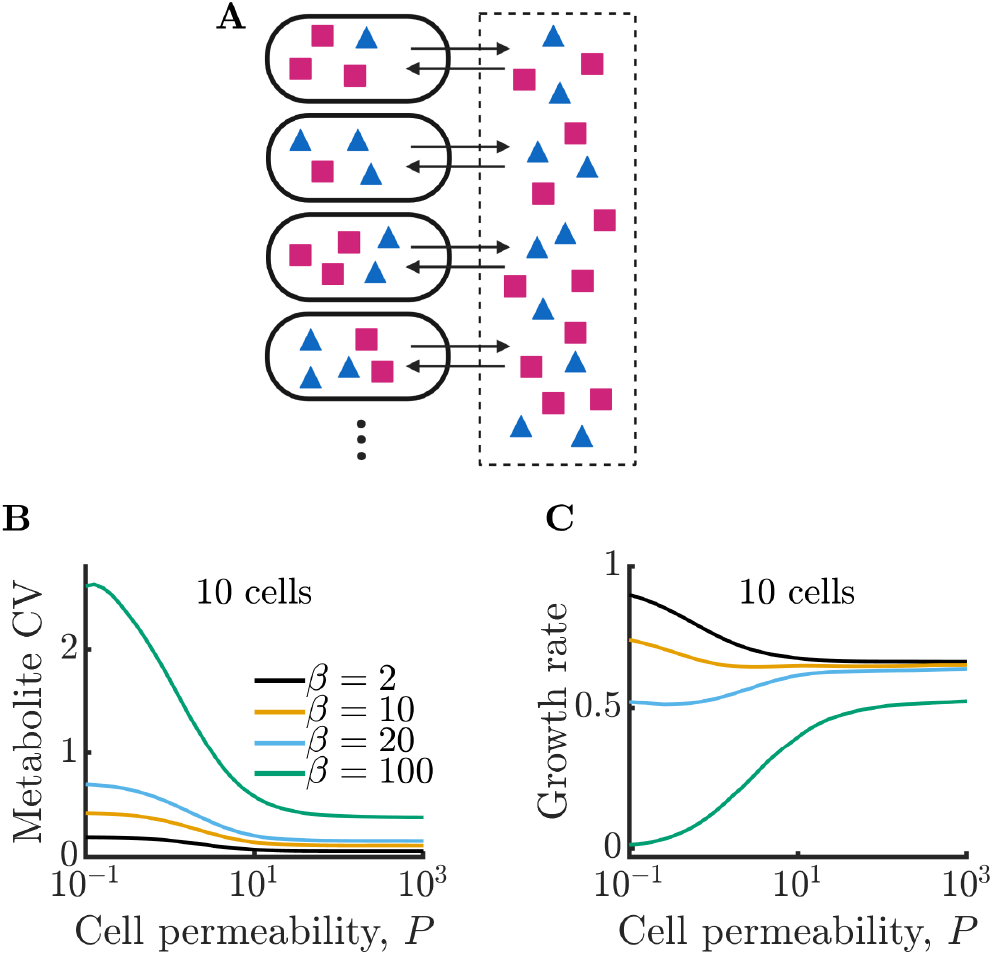
Bacterial cells can compensate for noisy enzyme regulation and increase growth rate by exchanging metabolites within a clonal community. (*A*) Schematic of multi-cell metabolism model. Individual cells regulate their own enzyme levels, but metabolites leak into the local medium and can be used by other cells in the community. (*B*) Intracellular metabolite coefficient of variation (CV) for a community of 10 cells as a function of cell permeability. (*C*) Average growth rate for community of 10 cells as a function of cell permeability. Parameters in *B* and *C* same as in Figure *1E*.

Enzyme production within the cell is regulated based on internal metabolite levels, with the cell exclusively producing the enzyme corresponding to the currently least abundant metabolite. A flow-chart of this regulation scheme is shown in Figure 1B. To reflect the bursty nature of gene expression [25], we assume that enzymes are produced in Poisson distributed bursts with average size *β*. Cells produce enzymes at a rate proportional to their growth rate such that the rate of enzyme bursts is 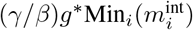, where *γ* is constant controlling the steady-state abundance of enzymes (such that at steady state 〈*E_i_*〉 = *γ/*2). Enzymes are diluted by growth at a rate proportional to their abundance 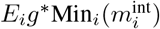. We model enzyme production as a stochastic process due to its intrinsically noisy nature, and model enzyme dilution as a deterministic process. The metabolite and enzyme equations are simulated numerically using a hybrid deterministic-stochastic method (see Appendix B for details).

In Figure 1C and D, we show example timecourses of metabolite dynamics for different burst sizes in cells with an average level of each enzyme of 〈*E_i_*〉 = *γ/*2 = 25. In Figure 1C, we show the metabolite levels of a cell with a small burst size *β* = 2. The small bursts allow for precise regulation of metabolite production, with the cell maintaining nearly equal levels of the two metabolites. In contrast, Figure 1D shows a cell with a burst size of *β* = 20. This cell’s poor enzyme regulation leads to imbalances in enzyme levels which in turn manifest as metabolite imbalances (see Appendix F —figure 4 for the corresponding enzyme timecourses).

**FIG. 3:**
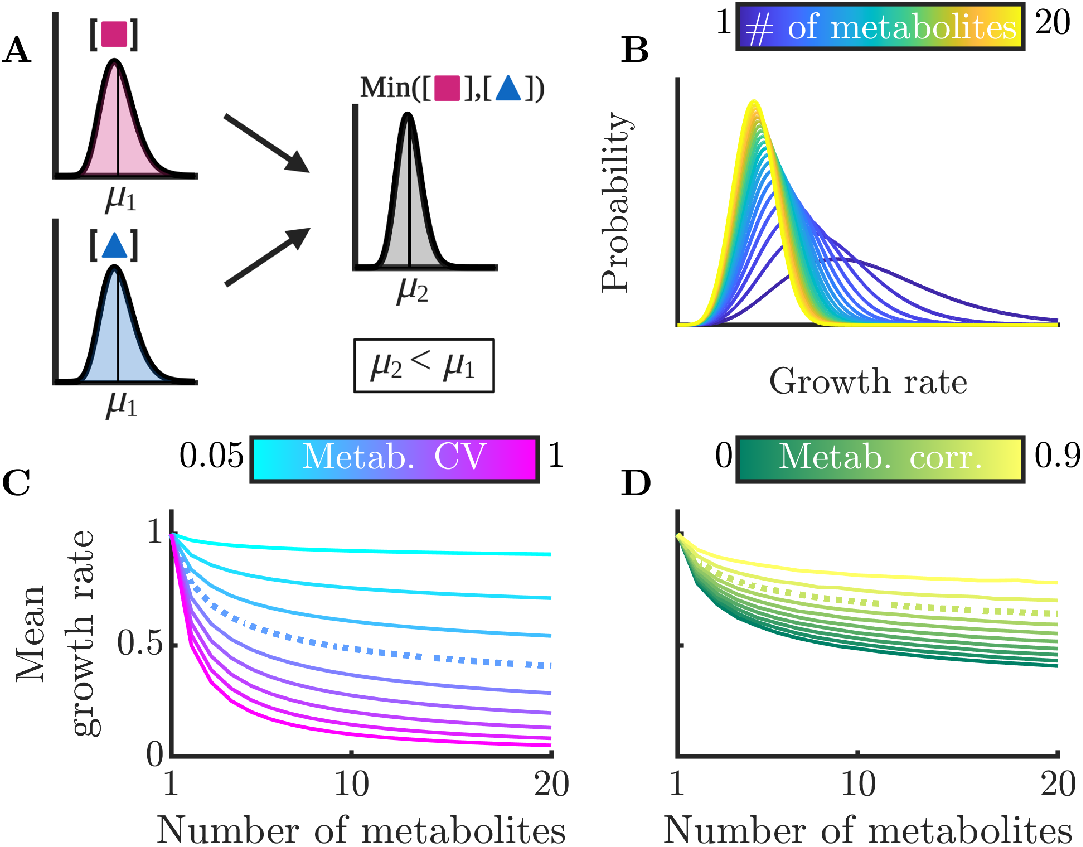
Sharing multiple metabolites can generically reduce noise and improve overall colony growth rate. (A) Implication of Liebig’s law of the minimum for fluctuating metabolites: the growth rate at any time is set by the lowest metabolite level, hence the average growth rate is lower than at the average metabolite level. The magnitude of this decrease grows with increasing metabolite variance. (B) Distributions of growth rates, set by minimum metabolite level Min(*m_i_*), for varying numbers of essential metabolites. (C) Mean growth rate as a function of number of metabolites and of metabolite CV. The dashed curve corresponds to CV = 0.4, as measured for essential proteins in *E. coli* [26]. (D) Mean growth rate as a function of number of metabolites and of the correlation coefficient between metabolites, for CV= 0.4. The dashed curve indicates a metabolite correlation of 0.7, approximately that observed for proteins in *E. coli* [26].

**FIG. 4:**
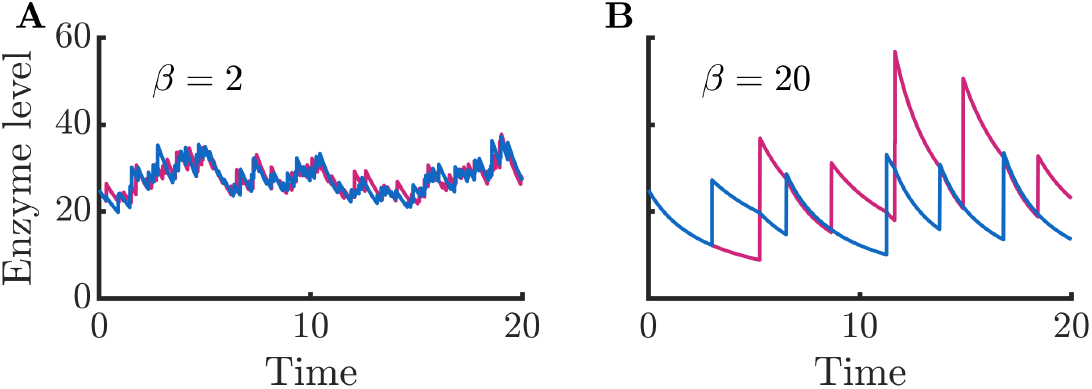
Enzyme timecourses corresponding to Figure 1CD. (*A*) Enzyme timecourse of a cell that produces enzymes in small bursts, same parameters as in Figure 1C. (*B*) Enzyme timecourse of a cell that produces enzymes in large bursts, same parameters as in Figure 1D. Note that the enzyme levels are substantially correlated with each other (*r* = 0.73 for *β* = 2 and *r* = 0.32 for *β* = 20).

How does the growth of an isolated cell depend on membrane permeability? In Figure 1E, we show the mean growth rate of isolated cells for varying permeability *P*. As can be seen, growth rate decreases monotonically with permeability. This follows because permeability leads to a loss of metabolites to the extracellular space where they cannot be utilized by the cell, but can be degraded. The coefficient of variation (CV) of the intracellular metabolite levels does not substantially change with increasing permeability, decreasing only very slightly due to stored metabolites in the extracellular space buffering fluctuations within the cell (see Appendix F—figure 5). Growth rate losses increase with growing extracellular volume, approaching the limit in which metabolites are permanently lost upon leakage from the cell.

**FIG. 5:**
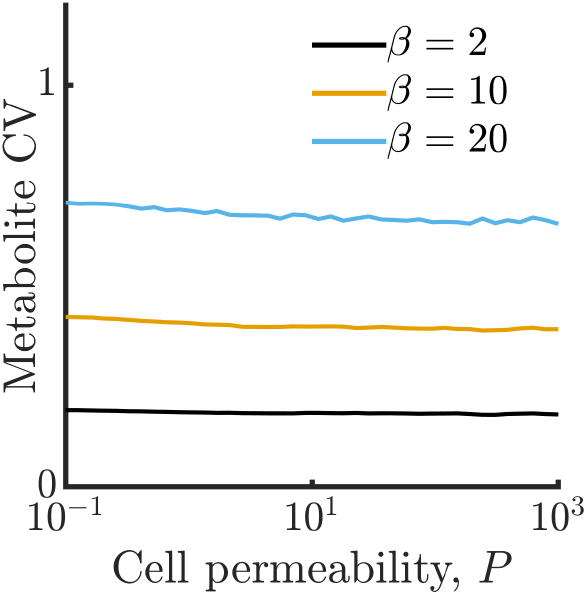
Metabolite CV corresponding to simulations in Figure 1E. All simulation parameters identical to those in Figure 1E. Data corresponding to *β* = 100 not shown as metabolite CV cannot be meaningfully estimated for cells with arrested metabolism.

The growth of isolated cells is also strongly influenced by the enzyme burst size, with small burst sizes permitting faster growth. This is seen in Figure 1F, which shows that growth rate decreases monotonically with average burst size *β*. The decreasing trend reflects a type of “use it or lose it” phenomenon in which cells grow poorly when they have large metabolite imbalances, as these result in metabolites being degraded instead of consumed for growth. The smaller the burst size, the lower the variance of the enzyme levels and the closer to equality metabolite production and levels can be maintained. Note that with sufficiently large burst sizes, the cell can experience irreversible metabolic arrest. This occurs because the cell must grow to produce additional enzymes, and if the cell experiences a sufficiently large metabolite imbalance, it may be unable to make another burst of enzyme before its existing metabolites are exhausted. In Figure 1E, this growth arrest is reflected in the *β* = 100 curve that is zero for all values of permeability. Similarly, in Figure 1F, there is a value of *β* beyond which cells do not grow.

### Multi-cell communities

We have characterized the behavior of an isolated model cell, but how do enzyme noise and metabolite leakage affect growth rates in a community? We now expand our model to a population of cells that share a common extracellular space with which they exchange metabolites, as depicted in Figure 2A. When cells leak there is now a possibility that these leaked metabolites will be taken up by other cells. To explore how leakage influences the collective metabolism of a multi-cell community, we simulate a community of 10 cells growing under the same conditions as in Figure 1E. In Figure 2B, we show the intracellular metabolite CV of these cells as a function of permeability for a range of enzyme burst sizes. As in the single-cell case, metabolite CV generally increases with increasing burst size. Interestingly, however, the metabolite CV now decreases substantially with increasing permeability. This occurs because metabolite exchange allows the cells to “average out” the noise arising from their individually poor enzyme regulation, a phenomenon we term noise-averaging cooperation (NAC). With large burst sizes, cells are prone to overproducing one type of enzyme, and thus overproducing a single type of metabolite. An isolated cell has no avenue to remedy this imbalance, leading to degradation of the overabundant metabolite. In a sufficiently large community this changes: within the population of cells, it is likely that there exist cells with opposite imbalances, and by exchanging metabolites these cells can collectively balance their metabolism.

How does this decrease in metabolite noise impact the average growth rate of cells within the community? In Figure 2C, we plot the growth rates of the communities from Figure 2B. In the case of a large burst size (*β* = 100), the improvement is extreme. With sufficiently high permeability, cells that were previously unable to grow at all due to their poor regulation can now grow at a substantial fraction of the optimal growth rate. We note that while metabolite CV decreases for all values of burst size, this does not always translate into a growth improvement. With very small burst sizes (*β* = 2), the increased permeability has the opposite effect and slightly decreases the growth rate. Since these cells already have efficient enzyme regulation, the moderate decrease in metabolite CV is outweighed by the increased degradation of metabolites within the extracellular space. Thus, the benefits of NAC are greatest under two conditions: 1) when individual cells have poor enzyme regulation and 2) when cells exist in a crowded space with minimal free volume between cells (such as in a biofilm).

How does community size influence the noise-reducing effect of NAC? To answer this question, we consider a simplified version of our model in which a population of *n* fully permeable cells are directly connected to each other. We track only a single type of metabolite and enzyme, assuming a constant growth rate such that the per-cell average rate of enzyme bursts is Γ/*β* and the rate of enzyme dilution is *μ_E_*. Metabolite consumption and degradation are aggregated into a single rate parameter *μ_m_*. For simplicity, we assume that all enzyme bursts are of size *β*, rather than being Poisson distributed. The Langevin equations for the total number of enzymes and total number of metabolites are therefore:

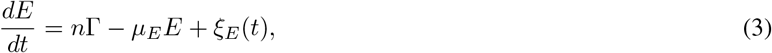

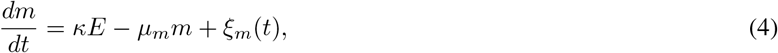

where the *ξ*(*t*) are noise terms with 〈*ξ*(*t*)〉 = 0 (see Methods for further details). Since these equations are linear, we can exactly compute the expression for the CV of the total enzyme level (see Appendix D):

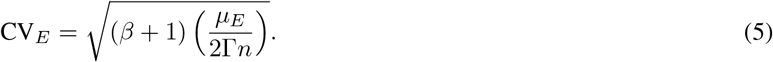

Consistent with our simulations, the CV increases with burst size *β*. The dependence on population size can also be immediately seen from this expression, with the CV being proportional to 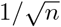. Thus, larger communities are expected to magnify the positive impact of metabolite exchange. We can also directly compute the metabolite CV:

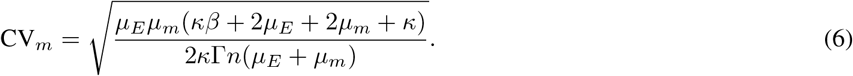

As expected, the metabolite CV has a similar scaling with respect to *n* and *β* as the enzyme CV.

These calculations, along with our simulations, characterize the potential benefits of NAC. The small size of bacteria make them inevitably noisy, possibly leading to growth losses. Metabolite leakage can act as a form of bacterial mutual aid, benefiting cells by allowing resource pooling.

### Generalized model framework

We have demonstrated NAC in a simple model with two metabolites, but how would the benefits apply to more realistic metabolic networks? We now develop a general framework to determine the impact of enzyme fluctuations and metabolite sharing on the growth of bacteria with an arbitrary number of non-substitutable metabolites. We begin with an arbitrary probability distribution function (PDF) of the intracellular levels of an individual metabolite *f* (*m_i_*). For simplicity, we first consider the case of independent but otherwise identical metabolites. For a given number of non-substitutable metabolites, we can then apply Liebig’s law of the minimum to compute the distribution of growth rates *q_n_*(*g*) by determining the distribution of the lowest metabolite level among a set of *n* metabolites. Note that while this minimum is technically only proportional to growth rate, for brevity we assume *g*^∗^ = 1 such that *g* = Min_*i*_(*m_i_*). To calculate *q_n_*(*g*), we use the cumulative distribution functions (CDFs) *F* (*m_i_*) and *Q_n_*(*g*):

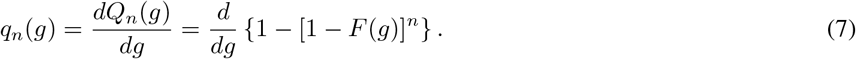

Intuitively, the mean of the distribution of growth rates with *n* > 1 non-substitutable metabolites will always be lower than the mean of *m_i_*, as shown schematically in Figure 3A. Thus, as in our simple two-metabolite model, the mean growth rate will depend not just on the means of the individual metabolite counts, but also their variability.

To examine this general model in more detail, we now use realistic metabolite distributions to quantitatively analyze the effect of various factors on the growth-rate distribution. While the distribution of metabolite levels in single cells has not been directly measured, there have been extensive measurements of single-cell protein distributions showing that these levels are typically gamma distributed. As we expect that enzyme levels are the dominant source of metabolite noise, we therefore approximate the metabolite distributions as gamma distributions, i.e. we assume that the metabolite distributions inherit the shape and thus the CV of the underlying enzyme distribution. In particular, we use the median gamma-distribution parameters measured for essential proteins in *E. coli* [26] as a base. We show the corresponding growth-rate distribution as a function of the number of non-substitutable metabolites in Figure 3B. One sees that as the number of metabolites increases, the mean of the growth-rate distribution decreases. This can be viewed as a ‘curse of dimensionality’: the more metabolites the cell must manage, the more likely it is that at least one will be poorly regulated at a given moment and constrain growth. Thus, NAC is most beneficial to cells that require large numbers of non-substitutable metabolites. In Figure 3C, we explore how this decrease in growth rate depends on the CV of the metabolite distribution, with the CV of the *E. coli* essential protein distribution shown as the dashed curve. The effect of the CV is to modulate the severity of the curse of dimensionality. If the metabolite levels are poorly controlled resulting in a large CV, the addition of more metabolites drastically reduces growth. Conversely, if the cell has tight control of its metabolites, it can manage significant numbers of non-substitutable metabolites without too large a growth loss. Interestingly, the curve corresponding to the CV of essential *E. coli* proteins shows a significant loss in growth rate, suggesting that real-world bacteria may suffer substantially from poor enzyme regulation. It should be noted, however, that enzyme count noise may overestimate noise in the resulting metabolites, as the enzymes can be regulated post-translationally, and metabolite fluxes may be buffered against enzyme fluctuations by network feedback effects [27].

Thus far we have assumed that the metabolites distributions are independent, but this likely does not hold in nature. Experimental studies have found that levels of different enzymes within the cell are correlated [26], suggesting that metabolite levels are likely also correlated. Indeed, this phenomenon occurs even in the simple models we analyzed above (see Appendix F — Figure 4). To account for these correlations, we computed the mean growth rate for varying degrees of correlation between metabolites. As a baseline, we again use the median distribution of essential proteins in *E. coli*. The results can be seen in Figure 3D, and show that correlation between metabolites reduces the adverse effects of metabolite noise. This occurs because if the metabolite levels are correlated, it is less likely that there will be a single outlying low metabolite level constraining growth. The correlation between certain proteins in *E. coli* has been measured [26], and we show this value as a dashed curve. While this level of metabolite correlation does improve growth, the growth loss associated with realistic enzyme noise is still substantial.

What if the growth rate is not determined by Liebig’s law of the minimum? Real growth functions are unlikely to be quite so simple, and given the variation that exists in microbial metabolism, it is unlikely that there is a single universally applicable growth function. Despite this uncertainty, we can determine what classes of growth function will lead to noise-driven growth defects. Consider an arbitrary growth function *g*(*X*) and vector of randomly varying metabolites *X*. We can express the decrease in growth due to noise as 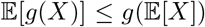. This statement is equivalent to the multivariate Jensen’s Inequality for concave functions, meaning that if *g*(*X*) is concave, the introduction of metabolite noise will decrease the mean growth rate. Growth functions in which the benefit of increasing individual non-substitutable metabolite levels is saturating will generally be concave. Thus, most reasonable growth functions will lead to decreased mean growth in the presence of metabolite noise. To demonstrate this, in Appendix F - figure 7 we show a version of Figure 3C with an alternative growth function based on the rate of protein synthesis. Interestingly, the dependence of the noise-drive growth loss on the concavity of *g*(*X*) implies that the magnitude of the loss may depend on the mean metabolite levels. If the metabolite levels are well above the saturation point of the growth process, such that the local growth function has low concavity, there will be minimal growth losses due to metabolic noise. A similar reduction in growth losses may occur if the metabolites are far below saturation.

**FIG. 6:**
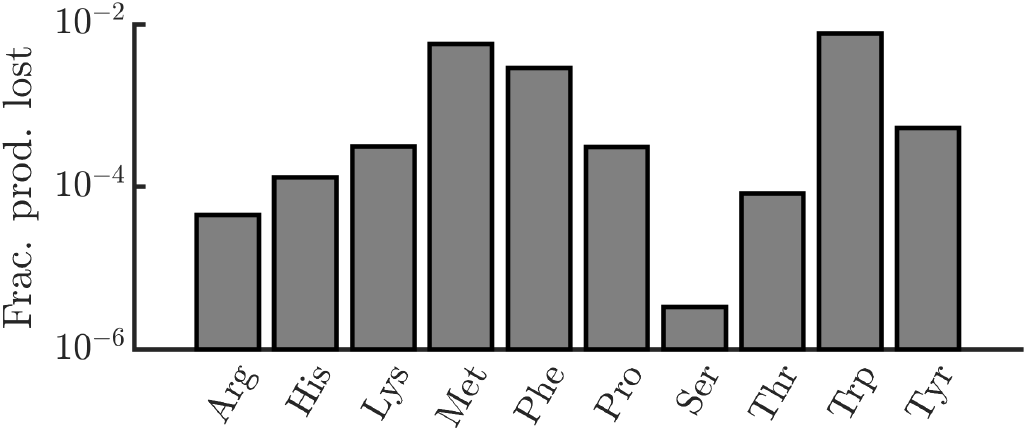
Estimated fractional production losses of amino acids in *E. coli* due to leakage. In order to determine the significance of amino-acid leakage in *E. coli*, we estimated the fraction of *E. coli*’s amino-acid production that is lost to leakage. Formally, we define the fraction of production lost as the ratio of the number of amino acids lost via leakage over the period of one division to the number of amino acids required to produce a daughter cell. For each amino acid, we require three experimentally measured quantities for this calculation: 1) the rate of leakage through the cell membrane, 2) the intracellular concentration of the amino acid, and 3) the number of amino acids required to produce a daughter cell. For leakage rates, we use data from artificial liposomes [23]. In cases where multiple pH conditions were tested, we used data measured at pH 7 (though leakage rates did not vary substantially with pH). This study measured data for only a limited set of amino acids. For other amino acids, we estimated their leakage rates using a linear regression of leakage rate versus log octanol/water partition coefficient (*r*^2^ = 0.96). Leakage rates were also adjusted for the differing size of the liposomes and typical *E. coli* cells, assuming a liposome radius of 200nm and an *E. coli* radius of 400nm [48]. Intracellular concentrations were taken from [24] and per-cell pool sizes were calculated assuming a cell volume of 1.8 10^−15^*L* [49]. A cell’s amino-acid production was assumed to be the number of amino acids required to produce a daughter cell, taken from [34]. With all of these experimental values, the fraction of production lost is 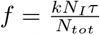 where *k* is the leakage rate, *N_I_* is the intracellular molecule count, *τ* is the doubling time (assumed to be 24 minutes), and *N_tot_* is the number of amino acids required to produce a daughter cell.

**FIG. 7:**
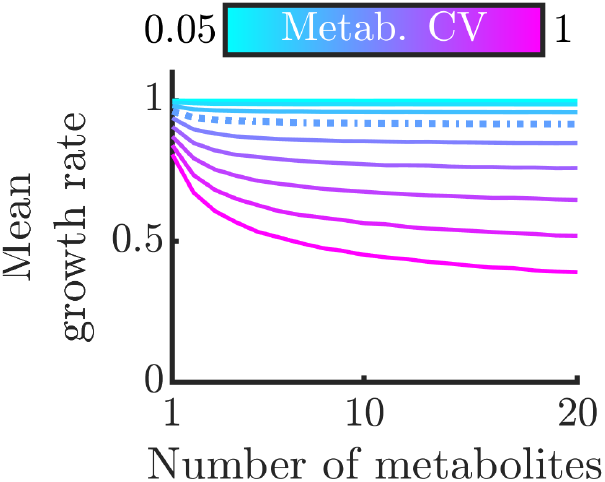
Version of Figure 3C with an alternative growth function. To demonstrate that our findings are not limited to Liebig’s law of the minimum, we repeat the analysis in Figure 3C with an alternative growth function from [50]. The function is 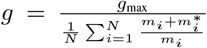, where *g*_max_ is the maximum growth rate, *N* is the total number of unique metabolites, and 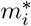 is the half-substrate constant of each metabolite. In this analysis we assume *g*_max_ = 1 and 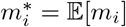.

With the above generalized framework, we were able to incorporate experimental measurements into our theory. Our preliminary analyses based on enzyme level measurements suggest that real bacteria may indeed suffer from substantial noise-driven growth defects. Combined with our analyses of simple metabolic models, this raises the possibility that bacteria can engage in NAC to improve their collective growth rate, particularly in tightly-packed environments like biofilms.

## Discussion

In this work, we develop a theory of noise-averaging cooperation (NAC), a novel mechanism potentially underlying the evolution of both intraspecies cooperation and interspecies cross-feeding. NAC allows microbes with individually poor regulation to average out their metabolic noise and raise their collective growth rate by sharing metabolites. Since NAC is strongest in crowded environments, it suggests an additional benefit of the biofilm mode of growth. With respect to cross-feeding evolution, our mechanism can be viewed as a generalization of the Black Queen Hypothesis: it provides a mechanistic explanation for the evolution of metabolite leakage, setting the stage for the emergence of metabolic interdependencies via gene deletions [28]. However, while we have shown that this mechanism can occur under plausible assumptions about metabolite noise and is consistent with some existing data, further study is needed to determine whether NAC occurs in nature and to what extent it may be a driver of cross-feeding.

How can we identify NAC in natural systems? Our theory predicts, counterintuitively, that it may benefit organisms to secrete essential metabolites into the environment. Thus, deliberate leakage or export of essential metabolites, such as amino acids or vitamins, is a potential signature of NAC. The clearest signature of deliberate export would be the existence of dedicated transporters for essential metabolites. Deliberate export could also occur via membrane leakage, but this case is more ambiguous as it is difficult to determine whether such leakage is “deliberate”, i.e. allowed by the cells, or is an unavoidable consequence of membrane permeability. As a test case, we examined the export of amino acids, a key class of non-substitutable metabolites, in *E. coli*. How much of *E. coli*’s amino-acid production is lost to leakage? Using prior experimental measurements of membrane permeability, intracellular concentrations, and amino-acid production rates, we estimate that *E. coli* loses only a small fraction (*<* 1%) of its amino-acid production to membrane leakage (see Appendix F — figure 6). This suggests that membrane leakage is not a substantial avenue of amino-acid export in *E. coli*. On the other hand, there exist multiple amino-acid exporters in *E. coli* [29–33], implying that *E. coli* does indeed engage in deliberate export of essential metabolites, consistent with NAC. Moreover, consistent with the idea that these exporters enable metabolic exchange, it has been shown that artificial auxotrophic *E. coli* strains can indeed cross-feed each other amino acids [34]. Note, however, that there may be reasons other than NAC for deliberate export of amino acids, such as overflow metabolism or the use of amino acids as signaling molecules [29, 35, 36].

Another way to probe the relevance of NAC to real bacteria would be to obtain more accurate estimates of intracellular metabolite distributions. Due to a lack of direct measurements of metabolite concentrations within single bacterial cells, we approximated the metabolite distribution using data from single-cell proteomics measurements. This approximation allowed us to estimate growth loss due to noise, but our conclusions depend on the assumption that the magnitude of metabolite noise roughly follows that of enzyme noise. In our simple metabolic model, this assumption is borne out, as the dominant source of metabolite noise is the burstiness of the enzyme dynamics. However, in real cells there are likely additional regulatory feedbacks that suppress metabolite noise. Thus, our analysis may overestimate metabolite noise and thus the benefits of NAC. Future theoretical studies could yield a more accurate estimate of metabolite noise using realistic models of intracellular metabolite dynamics that incorporate complete pathways and phenomena such as post-translational regulation. There is also the possibility that direct measurements of intracellular metabolite distributions will become available, as the technology for single-cell metabolite measurement is rapidly advancing [37, 38].

In addition to more precisely quantifying intracellular metabolite noise, better predicting noise-driven growth losses will require understanding the relationship between individual metabolite levels and growth rate in real cells. We showed that in order for metabolic noise to decrease mean growth rate, the only requirement is that the growth function be concave. We expect most growth functions to meet this condition, as the benefit of increasing metabolite levels generally saturates. However, it is possible that cells mitigate the impact of noise by maintaining their metabolite levels in a region of the growth function with low concavity, e.g. in a linear regime or near saturation. Thus, experimental data is needed to determine how sensitive cell growth is to metabolic noise. Experimental quantification of growth rate functions will likely require simultaneous measurement of growth and intracellular metabolite levels in single cells. A possible experimental system is a ‘mother machine’ [39] in which the growth rate of cells, and ideally metabolite levels, can be accurately tracked over long times. The technology for real-time single-cell metabolite measurement is rapidly developing: for example, a fluorescent reporter for branched-chain amino acids has recently been demonstrated in eukaryotes [38].

Even if the growth loss due to metabolic noise is large, for NAC to be beneficial the reduction in noise must outweigh the cost of metabolite loss in the extracellular space. In the context of a biofilm, some of this loss will likely arise from diffusion away from the biofilm, and thus the loss rate can potentially be calculated from the geometry of the biofilm and the diffusion constant of the metabolite within the biofilm matrix. Estimating the impact of other, spontaneous or reaction-based, forms of metabolite loss will likely require experimental measurements.

Analysis of intracellular metabolite dynamics and realistic growth functions may provide some support for NAC, but definitive testing of the mechanism will likely require dedicated experiments. *E. coli* would be a suitable organism for such experiments, as it is known to encode transporters for at least some essential metabolites, and has already been shown to engage in intercellular exchange of amino acids [34]. To directly test the theory, experiments will require at least two conditions: one in which cells are isolated and another in which they exist at a relatively high density. One possibility is to compare planktonic and biofilm cells, while another would be to assemble varying densities of planktonic cells. With isolated and crowded conditions defined, there are two major predictions that could be tested: growth rate should increase when cells are in crowded environments, and metabolic noise should be reduced when cells are in crowded environments.

Testing of the growth-rate prediction could be performed using population-level measurements of well-mixed cultures. The simplest way to test this would be measure the exponential-phase growth rates of bacterial cultures of different densities. An exponential-phase culture of *E. coli* could be resuspended at different densities in minimal media with saturating concentrations of nutrients, and growth rates measured (e.g. via OD). If NAC is occurring, the growth rate should be positively correlated with the culture density. Another possible experimental system is an *E. coli* chemostat fed with minimal media. The measured output in this system will be the steady-state biomass. NAC predicts that, compared to the case where growth rate is independent of density, the cell density will be higher than expected at low dilution rates (see Appendix E for details). For both of the above experiments, it will be important to determine whether the observed growth differences are due to metabolite exchange. To test this, one could employ mutants with different essential metabolite exporter genes deleted and measure whether the difference in growth rate still exists between isolated and dense conditions. Data from these mutants will need to be interpreted carefully, e.g. due to redundant/undiscovered transporters or unintended effects of the deletions.

Testing whether metabolic noise decreases with cell density will require techniques with single-cell resolution. The difficulty in testing this prediction will stem from finding a method to measure single-cell metabolite concentrations. One possible method to estimate the metabolite distribution is to measure timeseries of metabolite levels using the fluorescent reporter approach discussed above. One could image these fluorescent reporters in two-dimensional colonies and attempt to correlate the observed metabolic noise to local cell density. Similar to the earlier proposed experiments, one could employ export mutants to determine whether observed decreases in noise are due to metabolite exchange.

If NAC does exist in nature, how is it implemented and regulated by cells? It is unlikely that cells would exchange their entire metabolome with the external environment, but how would cells select which metabolites to exchange? If the metabolites within cells have different levels of noise, it might be optimal for cells to engage in NAC with the metabolites with the highest noise. This would imply that metabolites produced in small quantities are good candidates for NAC. The choice of which metabolites to exchange is also influenced by the architecture of the cell’s metabolic network. If there is a bottleneck in the network, for example if catabolism generates a small set of metabolites that are then used in a myriad of anabolic synthesis processes, it may be beneficial to exchange these bottleneck metabolites. In addition to selecting which metabolites to exchange, there is also the issue of deciding when to engage in NAC. It would be harmful for cells to engage in NAC at low cell density, as most of the secreted metabolites would be lost. Thus, regulation of NAC would likely be linked to quorum sensing. Note, however, that sensing of kin cell densities would not be sufficient for regulation of NAC. If the environment also contains a high density of non-kin, there will likely be a high effective loss rate of metabolites to these non-kin, making NAC disadvantageous. Thus, regulation of NAC should be dependent on multiple quorum-sensing circuits, using both kin and non-kin autoinducers [40].

We have thus far considered NAC in well-mixed environments, but real microbial communities can be highly spatially structured, and this has been shown to give rise to a number of behaviors not seen in well-mixed systems [41, 42]. How might NAC be affected by spatial structure? One consequence of spatial structure is the formation of nutrient gradients within bacterial biofilms, with bacteria nearing the nutrient source experiencing higher nutrient concentrations [43]. As a result, some cells in a biofilm will have higher metabolite production rates than others, and it may or may not be collectively beneficial for cells in high-nutrient areas to share their metabolites with those in low nutrient areas. In such situations, the diffusion coefficient of the shared metabolite will play a significant role, as this will determine how widely the metabolites are shared. In addition to spatial structure, fluid flow within environments may also impact NAC. For example, consider a microbial community growing in pipe-flow conditions. Cells at the beginning of the flow will have little incentive to engage in NAC, as all their resources will be lost to flow. Conversely, cells further downstream would benefit from NAC as they receive metabolites from their upstream neighbors. Given the ubiquity of spatial structure and flow in real microbial communities, probing the impact of these factors on NAC is a promising direction of future study.

We have focused on bacteria in this manuscript, but it is also possible that NAC may be relevant in other domains of life. For example, NAC highlights a potential advantage of multicellularity: a multicellular tissue separated from its external environment is the optimal environment for NAC. NAC may also apply to macroecological systems if the outcome of foraging for non-substitutable resources (such as food and water) is highly variable. Under such conditions, it may be beneficial for animals to engage in resource sharing, potentially supporting the development of social groups.

While much research is needed to determine the relevance of NAC to real bacteria, the theory highlights an interesting aspect of ecology: noise at even the smallest scales can have a dramatic impact on the entire ecosystem. In this manuscript we have focused on the single-species case, but there is potential for more novel behaviors in the many-species context, as has been observed in other resource-competition models [44]. Overall, we hope this work can serve as a foundation for further theoretical and experimental work on how noise in resource acquisition impacts ecology and evolution.

## Acknowledgments

We thank Matt Black for useful discussions on possible experimental tests of the theory. JGL was supported by an NSF GRFP (DGE-1656466). This work was supported in part by the National Science Foundation, through the Center for the Physics of Biological Function (PHY-1734030).

## I. METHODS

All code and data used in this manuscript can be found at https://github.com/jaimegelopez/NAC. Details on individual analyses and derivations can be found in the relevant appendices.

## Appendix A: Nondimensionalization of the two-metabolite model

To show how we obtained the nondimensionalized Equations 1–2, we begin with a generalized set of equations in which a parameter *α* relates the minimum of the two metabolite levels to the consumption rate of the metabolites:

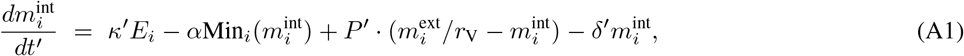

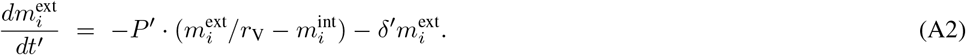

Here, 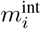 is the intracellular count of metabolite *i*, *t*′ is time, *κ*′ relates enzyme count to metabolite production, *E_i_* is the count of enzyme *i*, *α* relates the minimum of metabolite levels to metabolite consumption, *P*′ is permeability, 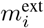 is the extracellular count of metabolite *i*, *r*_V_ is the ratio of extracellular to intracellular volume, and *δ*′ is the metabolite degradation rate. We now show that by rescaling time, we can eliminate *α*. We first introduce our dimensionless time variable *t*′ = *tu*, where *u* is a parameter with units of time. Substituting into the above equations yields:

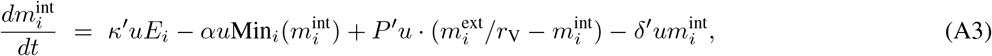

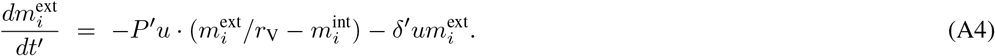

By defining *u* = 1/*α*, *κ* = *κ*′/*α*, *P* = *P*′/*α*, and *δ* = *δ*′/*α*, we recover Equations 1–2.

We present transport in our model as passive exchange with the environment, but these equations describe a broader range of transport phenomenon. In particular, our expressions for metabolite exchange are mathematically equivalent to active transport in the linear regime. To demonstrate this, suppose that import and export are mediated by two separate enzymes with different kinetics. In the linear regime, the transport expression will be 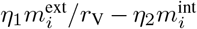 where the *η* are constants defining transporter performance (for Michaelis-Menten kinetics, each *η* would be the ratio of the maximum rate and the half-saturation constant of the corresponding transporter). Rearranging terms, one obtains 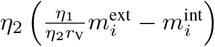 such that there is an effective permeability *η*_2_ and effective volume ratio 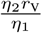.

## Appendix B: Hybrid numerical methods

In our two-metabolite model, the enzymes are produced in intermittent bursts which must be modeled stochastically, while other reactions in the system (such as metabolite consumption) involve a large enough number of molecules or occur frequently enough that they can be treated deterministically. In order to efficiently simulate this system, we use a hybrid stochastic-deterministic algorithm that can be viewed as a generalization of the Gillespie algorithm. We briefly present the rationale for the algorithm here, with a full description available in [45].

We begin with the traditional Gillespie algorithm. In this algorithm, the time between events is an exponential random variable with mean 1/*r*_tot_, where *r*_tot_ is the sum of all reaction rates in the system. This formulation assumes that *r*_tot_ is constant between stochastic reactions, something that will not be true in a system that also includes deterministic reactions. To account for time-varying reaction rates, we first recognize that the Gillespie event condition can be rewritten as:

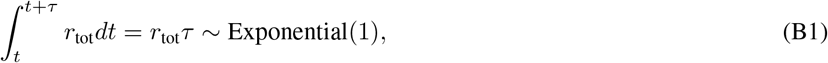

where *τ* is the stochastic time before the next reaction. This rewritten condition makes intuitive sense: if one regards the integral on the left-hand side as a kind of cumulative probability, this states that for every one unit of cumulative probability, on average one event occurs. From Equation B1 we can rigorously construct a simulation algorithm for systems with time-varying reaction rates. We first sample a random number *x*_1_ ~ Exponential(1). We then integrate the deterministic dynamics until we find a *τ* such that:

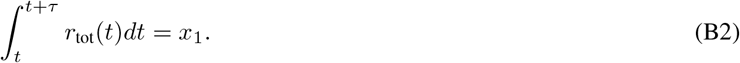

In our method, the integration is done by Euler’s method. We then sample another random number *x*_2_ ~ Uniform(0, 1) and use this to determine which reaction occurs, as in the traditional Gillespie method. The reaction event then occurs at time *t* + *τ*, and we repeat the above procedure until the simulation reaches a termination condition. In our simulation, the only stochastic events are the enzyme production bursts, while all other reactions are modeled deterministically. For calculation of CVs and growth rates in Figures 1 and 2, simulations were run for 4 × 10^5^ time units, with statistics computed from the last 3 × 10^5^ time units. For Figure 1EF, 140 replicate simulations were run per point. For Figure 2BC, 40 replicate simulations were run for most points, with 200 replicates run for certain simulations with large burst size and low permeability.

## Appendix C: Generalized framework

To compute distributions of growth rates from metabolite distributions, we first compute the growth distribution’s CDF in accord with Equation 7 and then apply numerical differentiation to yield the PDF. To obtain the expected value, we numerically integrate according to the expression 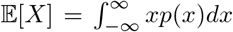 where *p*(*x*) is the probability density of value *x*. As a reference distribution, we used the median gamma distribution of *E. coli* essential proteins observed in [26] (*k* = 6.4 and *θ* = 5.2, taken from table S3). For distributions with different CVs, we maintain the same mean as the median distribution of *E. coli* essential proteins.

The case of correlated metabolite levels was analyzed by simulation. To generate correlated distributions, we first generate 50,000 samples for each metabolite from a multivariate normal distribution with the desired correlation structure. The MATLAB function mvnrnd is used for this purpose. We then use the method of copulas to generate samples from correlated gamma distributions [46]. This method first involves transforming the normally distributed samples into uniformly distributed samples using the inverse CDF of the normal distribution. We then transform these uniformly distributed samples into gamma distributed samples using the appropriate gamma CDF. Note that this method is only guaranteed to exactly preserve the rank correlation, but we find that the linear correlation is also very well preserved.

## Appendix D: Langevin description

To better understand the influence of community size on metabolite noise, we consider a simplified version of the two-metabolite model. In this model, we only track the concentration of a single constitutively produced enzyme and its corresponding metabolite. The Langevin equations governing the dynamics are:

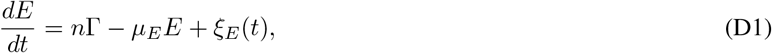

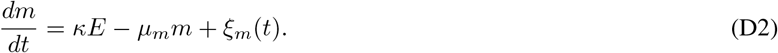

where the Langevin noise terms have zero noise 〈*ξ_E_*(*t*)〉 = 〈*ξ_m_*(*t*)〉 = 0 and are delta-correlated such that *ξ_i_*(*t*_1_)*, ξ_j_*(*t*_2_)〉 = Ω*_ij_δ*(*t*_1_ − *t*_2_), where *δ*(·) is the Dirac delta function. These equations have mean steady-state fixed point at 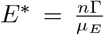 and 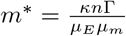. To analytically compute the CV of the enzyme and metabolite, we employ the method of Swain [47]. This method relies on linearizing the system about steady state and assumes that fluctuations about the steady state are sufficiently small so as not to drive the system out of the linear regime around the fixed point. Since our system is linear to begin with, this method will actually provide us with exact expressions of the moments.

We begin by computing the elements of Ω. The squared deviation of *ξ_E_*(*t*) will obey:

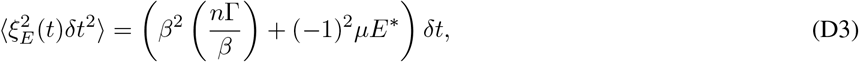

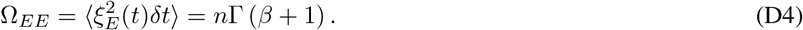

Similarly, we can compute the value of Ω_*mm*_:

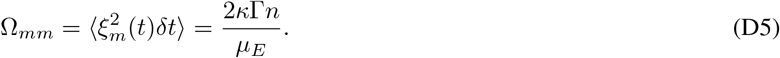

The off-diagonal entries of Ω will be zero, as the two equations share no common reactions.

Next, we represent our deterministic dynamics as a matrix equation by computing the Jacobian about the fixed point

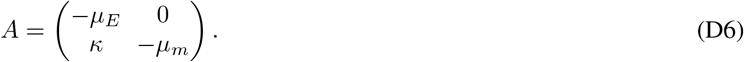

This Jacobian has eigenvalues *λ*_1_ = −*μ_E_* and *λ*_2_ = −*μ_m_* with a matrix of column eigenvectors

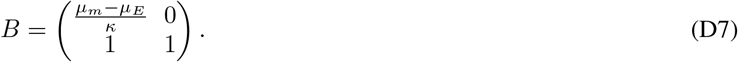

Variances and covariances of the state variables can then be computed using the following expression:

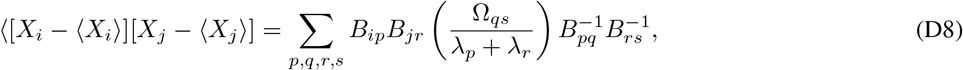

where the *X_i_* are the state variables.

Substituting in the values of *B*, Ω, and *λ* yields

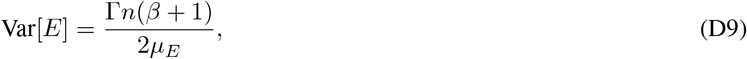

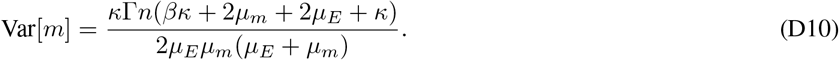

From these expressions for the variance, we compute the CVs as

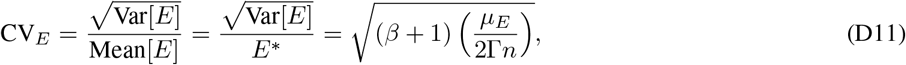

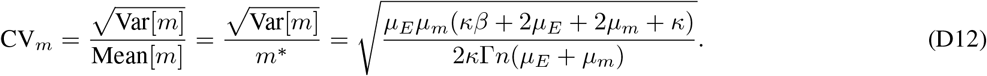

## Appendix E: Chemostat NAC experiment theory

Here, we present the theory for an experiment in which the effects of NAC are measured in a chemostat. First, we begin with the equations governing the chemostat dynamics:

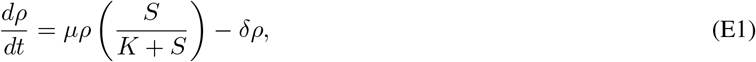

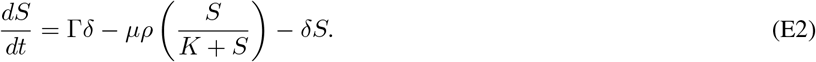

where *ρ* is the biomass concentration, *μ* is the maximum growth rate, *S* is the nutrient concentration, *K* is the half-saturation constant, *δ* is the dilution rate, and Γ is the inlet nutrient concentration. Note that we are measuring biomass and nutrients in the same units. The steady-state nutrient value will be:

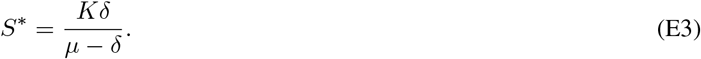

The steady-state biomass will therefore be:

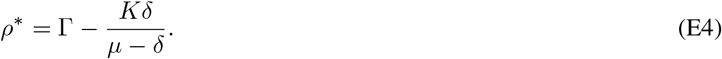

Thus, if we model NAC as an increase in *μ*, a higher than expected dilution rate will be required to maintain a given high cell density. To test this, one can estimate *μ* and *K* at low cell densities and compute the predicted density as a function of dilution rate using Equation E4. Then, the actual steady-state densities can be measured for a range of dilution rates. If NAC is occurring, the predictions will match the experimental data at high dilution rates, while there will be a significant discrepancy between predicted and observed densities at low dilution rates.

## Appendix F: Supplemental figures

In this section, we present supplemental figures that support the main text. Each figure’s caption contains all pertinent information.

